## Supplementary figures and images for "Global and local tension measurements in biomimetic skeletal muscle tissues reveals early mechanical homeostasis"

### Supplementary Figure 1_engineering drawing_bX

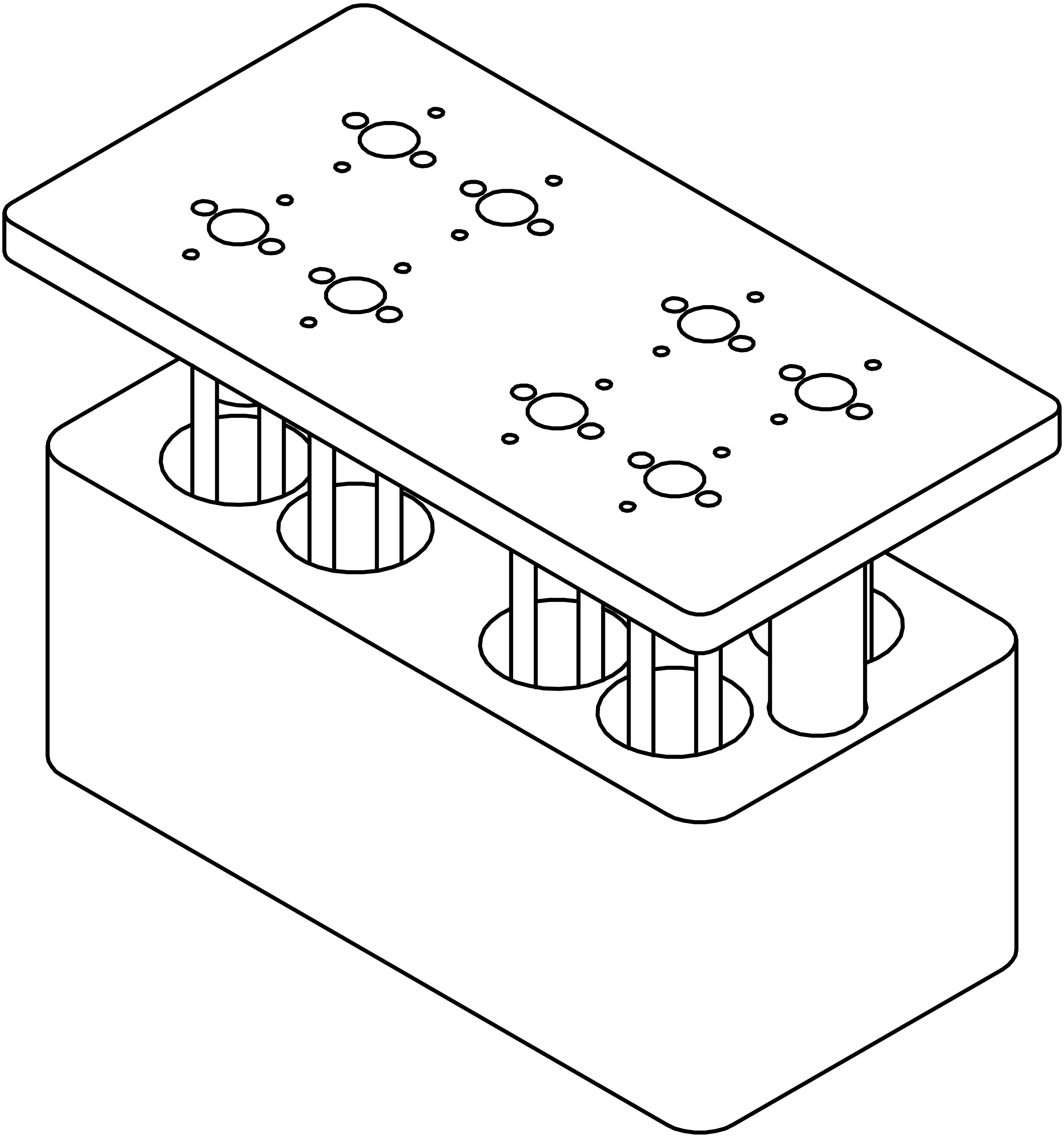

### Supplementary Figure 2 150kPa bead control

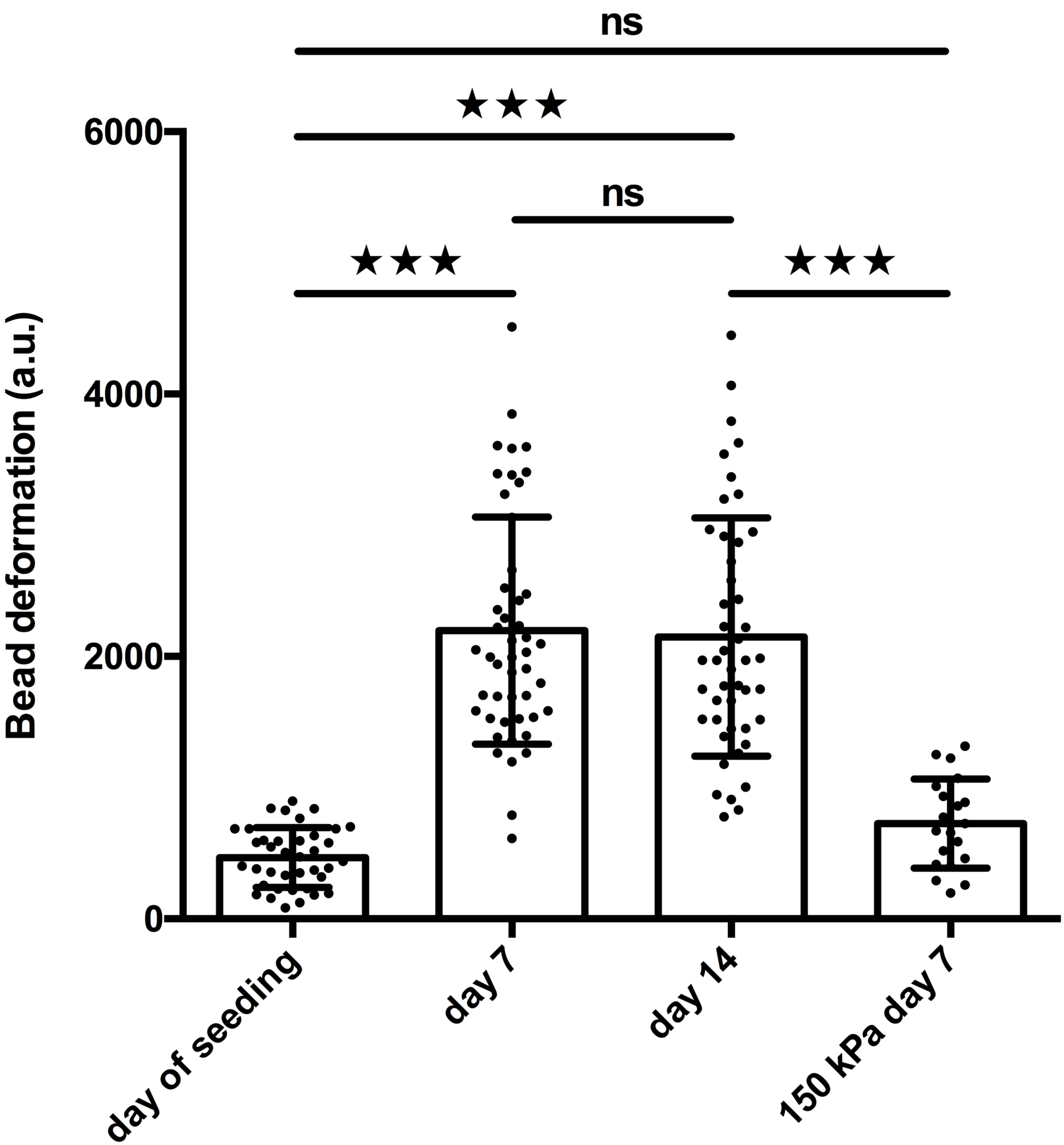

### Supplementary Figure 3 Fixation control

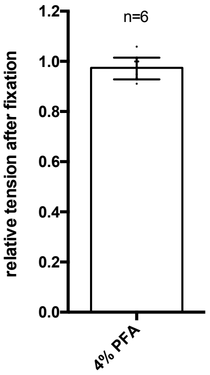
